# Developing Y chromosome sex ratio distorters in the model insect *Drosophila melanogaster*

**DOI:** 10.1101/2025.10.27.684751

**Authors:** Yael Arien, Chen Zacharia, Elad Yonah, Elad Bonda, Philippos Aris Papathanos

## Abstract

CRISPR-Cas9 sex ratio distortion (SRD) systems can suppress insect populations by biasing progeny toward males, but realizing such systems requires reliable Cas9 expression from insect Y chromosomes. Here, we tested whether the spermatocyte-specific β*Tub85D* promoter can drive functional Cas9 expression when inserted on the *Drosophila melanogaster* Y chromosome. Using CRISPR-mediated homology-directed repair, we generated a Y-linked β*Tub85D*-Cas9-T2A-eGFP construct and compared its activity with an autosomal counterpart. Whereas autosomal β*Tub85D*-Cas9 induced strong male-biased sex ratios when paired with an X-poisoning gRNA, the Y-linked construct failed to distort sex ratios and exhibited approximately 2,000-fold reduction in Cas9 transcript abundance. Nonetheless, weak but detectable GFP fluorescence and Cas9 transcripts confirmed partial Y-linked promoter activity. These findings provide the first direct experimental evidence of meiotic sex chromosome inactivation (MSCI) acting on the *Drosophila* Y chromosome, revealing that meiotic promoters can remain weakly active despite strong repression. This work defines transcriptional limits of the *Drosophila* Y chromosome and informs the design of next-generation Y-linked gene drives for sustainable insect control.

## Introduction

Genetic control of insect populations is a critical strategy for both modern agriculture and public health (Alphey 2014). Among the most promising approaches are sex ratio distortion (SRD) systems that exploit endonucleases like CRISPR–Cas9 to target X-linked sequences during male meiosis, either eliminating X-bearing gametes or causing lethality in female offspring (Galizi et al. 2016; Fasulo et al. 2020; Compton and Tu 2022; Haber et al. 2024). By producing an excess of males, these systems deplete the pool of reproductive females and can, in principle, lead to population suppression. While most SRD systems developed to date employ autosomal insertions as proof-of-concept demonstrations, these configurations are suboptimal for field application. Because autosomal transgenes are inherited by both sexes, they impose unnecessary fitness costs and create selection for resistance alleles in females. Restricting SRD components to the Y chromosome would ensure male-limited inheritance and activity, making such systems self-limiting and evolutionarily stable (Deredec et al. 2011; Burt and Deredec 2018; Simoni et al. 2020).

Several studies have demonstrated that engineered SRD constructs expressing endonucleases under the spermatocyte-specific β*2-tubulin* promoter can generate highly male-biased progeny through X-shredding or X-poisoning mechanisms when these are located on autosomes (Galizi et al. 2016; Fasulo et al. 2020; Meccariello et al. 2021; Haber et al. 2024). However, when such constructs were inserted onto the *Anopheles gambiae* Y chromosome, they failed to induce sex ratio distortion and exhibited no measurable transcription, indicating strong Y□linked silencing likely associated with meiotic sex□chromosome inactivation (MSCI; Alcalay et al., 2022).

Whether a similar barrier exists in *Drosophila* remains unknown (Fuller 1993; Vibranovski 2014). The *D. melanogaster* Y chromosome is largely heterochromatic and gene-poor but differs in composition and evolutionary history from that of *Anopheles* (Carvalho 2002; Krzywinski et al. 2006). Despite its heterochromatic composition, the *D. melanogaster* Y chromosome becomes transcriptionally active in primary spermatocytes, where giant fertility genes such as *kl-2, kl-3*, and *kl-5* are robustly expressed (Piergentili 2007). These genes form conspicuous, thread-like nuclear loops that extend into the nucleoplasm and correspond to regions enriched for RNA polymerase II and euchromatic histone marks (Bonaccorsi and Lohe 1991; Piergentili 2007). This unusual organization demonstrates that the *D. melanogaster* Y can support strong, stage-specific transcription in the male germline. However, it remains unclear whether exogenous transgenes can be expressed during meiosis from the *D. melanogaster* Y chromosome.

Unlike mammals, *D. melanogaster* males lack meiotic recombination and display a unique pattern of germline gene regulation. Whether its sex chromosomes undergo true meiotic inactivation is actively debated (Vibranovski 2014; Mahadevaraju et al. 2021).

Transcriptomic analyses consistently show that X-linked genes are underrepresented and less expressed in spermatocytes relative to autosomal genes (Vibranovski et al. 2009; Meiklejohn et al. 2011; Landeen et al. 2016). However, these patterns cannot be taken as direct proof of MSCI because several overlapping processes influence sex-chromosome transcription in the *D. melanogaster* germline. A central factor is dosage compensation, which equalizes X-linked expression between males (XY) and females (XX) in somatic tissues, and is thought to be generally absent in the male germline, causing lower baseline X-linked transcription regardless of meiotic silencing. Single□cell RNA□sequencing reveals limited pre□meiotic upregulation that disappears in spermatocytes (Witt et al. 2021; Anderson et al. 2023). Over evolutionary time, many testis-biased genes have relocated from the X to autosomes, while those remaining on the sex chromosomes may have evolved stronger or temporally shifted promoters that counteract repression (Vibranovski et al. 2009; Landeen et al. 2016). Consequently, genome-wide expression and epigenetic data conflate chromatin regulation with gene-specific adaptation and cannot by themselves reveal whether meiotic silencing operates on inserted transgenes.

Functional assays using transgenes offer the most direct test of meiotic transcriptional competence. When reporters driven by spermatocyte-specific promoters such as β*Tub85D* or *ocnus* are inserted on the X chromosome, expression drops several-fold relative to identical autosomal insertions (Hense et al. 2007; Kemkemer et al. 2011; 2014). These classic experiments demonstrate that the X-chromosomal context itself restricts transcription during meiosis, independent of gene evolution. By contrast, no comparable assay has tested whether a meiotically expressed transgene remains active when inserted into the Y chromosome of *Drosophila*. Previous Y-linked integrations in *Drosophila* have employed early germline promoters such as *vasa*, which drive Cas9 expression in premeiotic cells (Gamez et al. 2021). However, because these promoters are active before meiosis, they are not informative for evaluating whether the Y chromosome supports transcription of meiotically expressed transgenes that have not evolved compensatory mechanisms. Despite extensive use of meiotic promoters for autosomal transgenes, it is unknown whether such promoters can sustain transcriptional activity from the *D. melanogaster* Y chromosome. No study has directly tested whether a spermatocyte-specific transgene, such as *βTub85D*-Cas9, can overcome Y-linked chromatin repression to achieve functional meiotic expression. This knowledge gap limits both our understanding of Y chromosome regulation and the rational design of Y-linked gene drive systems.

We therefore engineered a *βTub85D-*driven Cas9–T2A–eGFP construct and targeted it to a defined locus on the Y chromosome using CRISPR-mediated homology-directed repair. We evaluated its ability to distort progeny sex ratios when combined with an X-poisoning gRNA targeting the haplolethal gene *RpS6*, and compared it with an autosomal *β2-tubulin*-Cas9 line. By coupling direct molecular assays with a functional SRD readout, this study provides the first direct evaluation of whether a meiotically expressed transgene can remain active when placed on the *Drosophila* Y chromosome. The results clarify the extent of Y-linked transcriptional competence during spermatogenesis, reveal key regulatory contrasts between *Drosophila* and *Anopheles*, and inform the design of Y-linked gene drive systems for sustainable insect population control.

## Results

### Generation of a Y chromosome *βTub85D*-Cas9 strain

To assess whether the *D. melanogaster* Y chromosome can support transcription of a spermatocyte-specific transgene, we generated a Cas9–T2A–eGFP cassette driven by the *βTub85D* promoter (hereafter *βTub85D*-Cas9) and inserted it into a defined Y chromosome locus using CRISPR-mediated homology-directed repair (HDR) **(Figure 1A)**. The construct was flanked by *gypsy* and CTCF insulators to mitigate heterochromatin-induced silencing and contained a 3xP3-tdTomato fluorescent marker for identification and selection of transformants. Homology arms corresponding to a previously validated intergenic Y-linked target site were incorporated to promote site-specific integration (Buchman and Akbari 2019; Gamez et al. 2021). We have previously shown that autosomal integrations of this transgene result in high levels of Cas9 activity, including strong male-biased sex ratios when combined with X-poisoning gRNAs targeting X-linked ribosomal protein genes (Fasulo et al. 2020; Haber et al. 2024).

**Figure 1.**
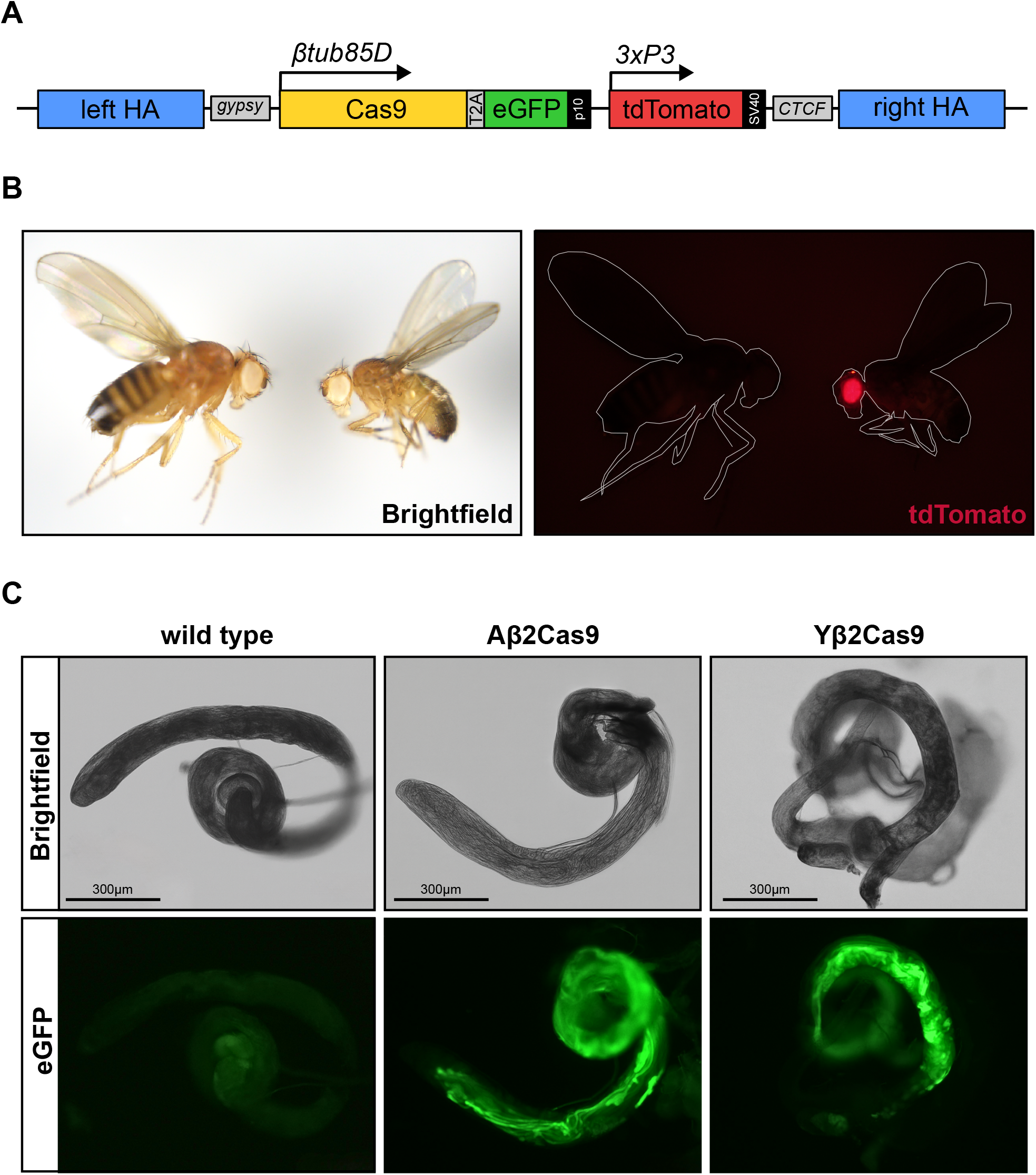
Generation of a Y-linked *βTub85D*-Cas9 strain. **(A)** Schematic of the *βTub85D*-Cas9 construct for Y chromosome knockin. Cas9 expression is driven by the spermatocyte-specific *βTub85D* promoter linked to eGFP with a T2A self-cleaving peptide. A 3xP3-tdTomato marker provides eye-specific fluorescence for transgenic selection. The transgene is flanked by *gypsy* and *CTCF* insulators and Y chromosome homology arms (left HA and right HA) for knockin using homology-directed repair. **(B)** Brightfield and tdTomato expression in male and female adults of Yβ2Cas9 strain. Eye-specific tdTomato fluorescence can be observed in males (right) but not females (left), confirming Y chromosome linkage. **(C)** eGFP expression in dissected testes from wild-type (WT), autosomal Cas9 (Aβ2Cas9), and Y-linked Cas9 (Yβ2Cas9) males. Top row: brightfield; bottom row: eGFP fluorescence. Scale bar = 300□μm.

As expected, transformation efficiency was low, consistent with prior reports of reduced HDR activity on the Y chromosome (Buchman and Akbari 2019; Gamez et al. 2021). From approximately 650 injected embryos, four F□ males exhibited tdTomato fluorescence in the eyes, three were sterile, and one successfully sired offspring. Eye fluorescence was subsequently observed exclusively in males of the progeny, confirming male-limited inheritance and stable Y-linkage of the transgene (**Figure 1B**; Table S1). The resulting line was designated Yβ2-Cas9.

Leveraging the T2A-eGFP marker, testes were dissected from Yβ2-Cas9 males to evaluate *βTub85D* promoter activity from the Y chromosome, and compared to those from an autosomal *βTub85D*-Cas9 line (Aβ2-Cas9) and wild-type controls. As expected, all Aβ2-Cas9 males displayed strong, uniform GFP fluorescence in the testes, reflecting robust spermatocyte-specific expression. In contrast, GFP was detected in only approximately 62 % of Yβ2-Cas9 males (**Figure 1C**), indicating that the *βTub85D* promoter retains partial activity when Y-linked but experiences variable or reduced transcriptional output.

### Evaluating sex ratio in progeny of Y-linked and autosomal *β-tubulin*-Cas9 males

To evaluate sex-ratio distortion (SRD) activity, we generated transheterozygous males carrying both the X-poisoning *RpS6_2* gRNA and either a Y-linked, an autosomal, or combined *βTub85D*-Cas9 construct. Each of these male genotypes was then crossed to wild-type females, and the sex ratios of the resulting progeny were scored to quantify SRD activity (**Figure 2**). Control crosses included Cas9-only males lacking the gRNA and wild-type intercrosses to account for baseline sex ratio variation **(Figure 3)**. Males carrying the autosomal construct (*Aβ2-Cas9/RpS6_2*) produced strongly male-biased progeny, with an average of 74.8 % ± 8.1 males, significantly higher than both Y-linked and Cas9-only controls (Tukey’s HSD, *p* < 0.01). Males carrying the Y-linked construct (*Yβ2-Cas9/RpS6_2*) produced nearly balanced progeny (53.8 % ± 6.2 males), not significantly different from their Cas9-only counterparts (*Yβ2-Cas9/+*, 47.7 % ± 6.2 males; *p* > 0.05). Interestingly, the strongest distortion was observed in males carrying both constructs (*Aβ2-Cas9+Yβ2-Cas9/RpS6*L), which yielded 85.8 % ± 4.9 male offspring, significantly exceeding all other groups (Tukey’s HSD, *p* < 0.001). Double-Cas9 males lacking the gRNA (*Aβ2-Cas9+Yβ2-Cas9/+*) displayed no sex bias (52.5 % ± 5.3 males; *p* > 0.05), confirming that the distortion required Cas9–gRNA activity **(Figure 3)**.

**Figure 2.**
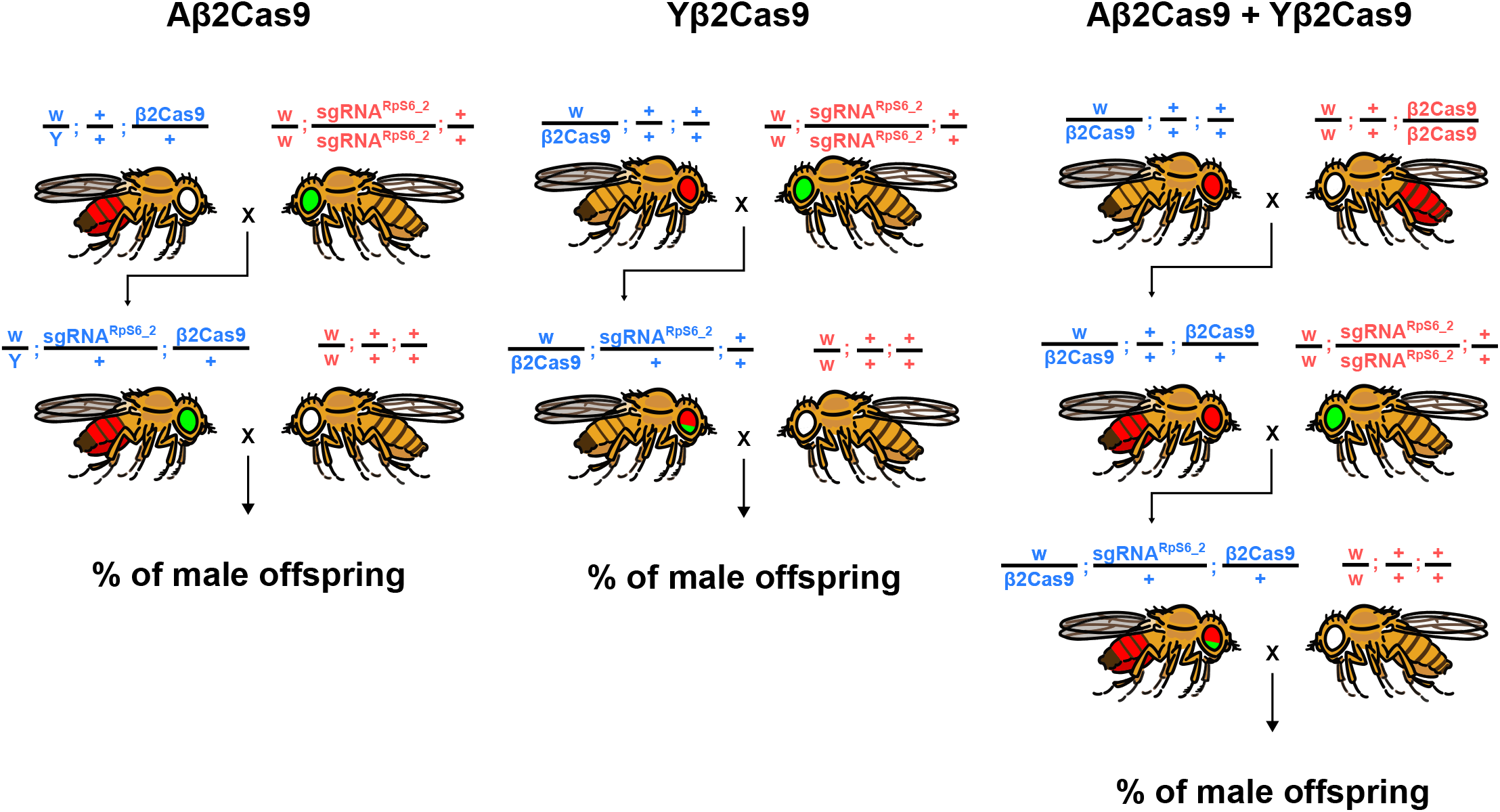
Experimental crosses used for sex ratio distortion assays. Crosses involved autosomal (Aβ2Cas9 - left panel), Y-linked (Yβ2Cas9 - middle panel), and dual (A + Yβ2Cas9 - right panel) Cas9 sources with an X-poisoning gRNA targeting the haplolethal X-linked gene *RpS6_2*. Aβ2Cas9 individuals are marked by an Opie2-DsRed reporter (indicated by red abdomens). Yβ2Cas9 individuals are marked by a 3xP3-tdTomato reporter (indicated by red eyes), and the sgRNA *RpS6_2* by a 3xP3-eGFP reporter marker (indicated by green eyes). F1 males from these crosses were used to assess sex-ratio distortion in their progeny.

**Figure 3.**
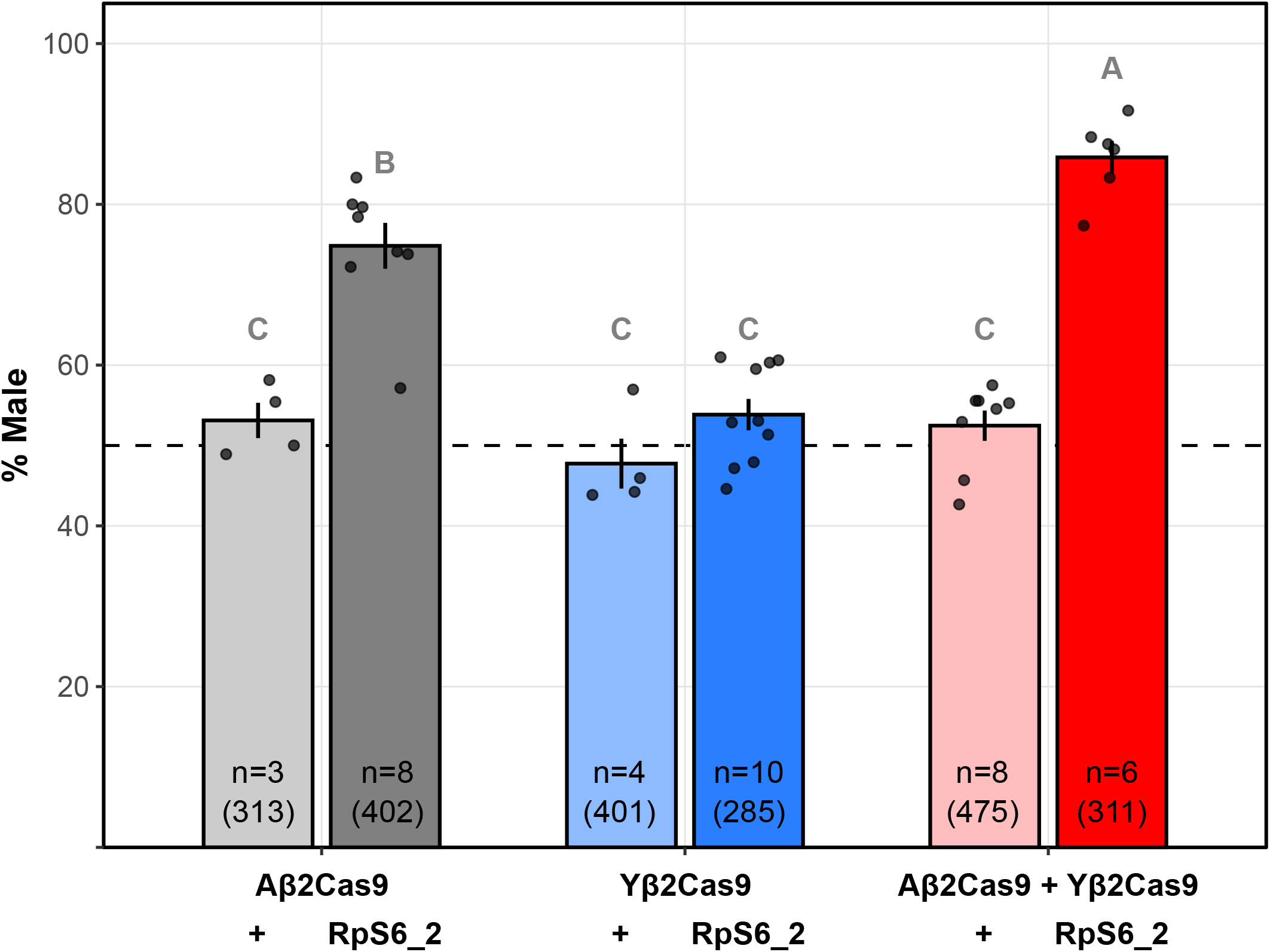
Cas9-driven sex-ratio distortion via targeting of the haplolethal gene *RpS6* during spermatogenesis. Bars represent the mean ± SE percentage of male progeny for each genotype. Autosomal βTub85D-Cas9 (Aβ2Cas9), Y-linked βTub85D-Cas9 (Yβ2Cas9), and the combination of both Cas9 sources (A+Yβ2Cas9) were each tested with or without the *RpS6_2*-targeting gRNA. Control males lacking the gRNA produced approximately equal sex ratios, whereas inclusion of the gRNA generated male-biased offspring, consistent with X-chromosome cleavage. The strongest bias was observed in the double-driver genotype (A+Yβ2Cas9/*RpS6_2*), indicating an additive effect of dual Cas9 sources. Letters above bars denote statistically distinct groups (Tukey’s HSD, *p* < 0.05). Numbers within bars indicate the number of replicate vials scored, with total progeny shown in parentheses.

**Figure 4.**
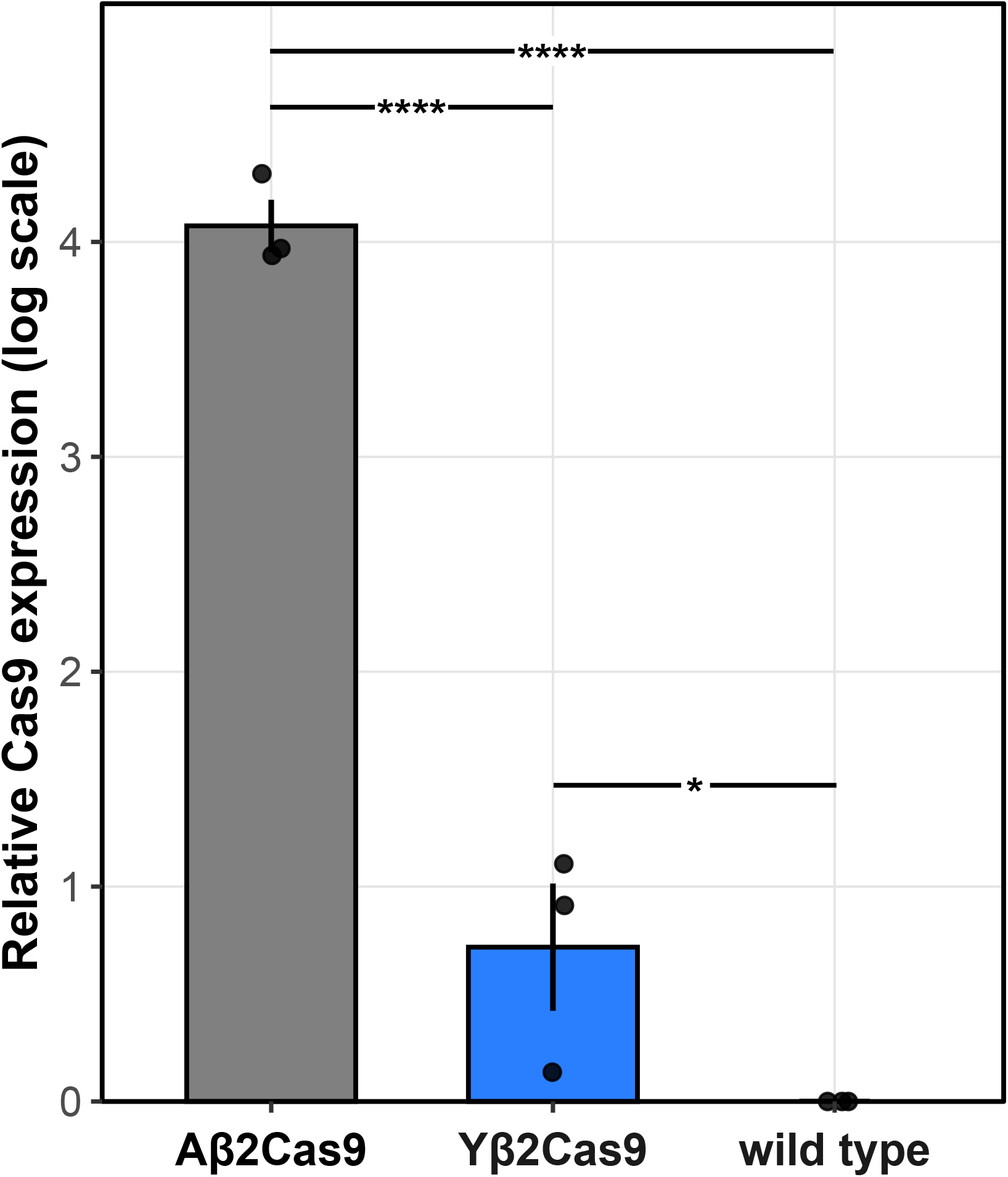
Relative Cas9 expression levels in testes of autosomal and Y-linked *βTub85D*-Cas9 lines. Bars represent the mean ± SD log-transformed fold change in *Cas9* transcript abundance, normalized to the reference gene *RpL32* and expressed relative to wild type (WT). Significantly higher *Cas9* expression was detected in autosomal *βTub85D*-Cas9 (Aβ2Cas9) males compared to Y-linked *βTub85D*-Cas9 (Yβ2Cas9) and WT controls. Statistical significance was assessed by one-way ANOVA followed by Tukey’s multiple comparisons test (*p* < 0.05; **** *p* < 0.0001).

Together, these results demonstrate that Y-linked Cas9 expression alone is insufficient to induce measurable SRD, while autosomal Cas9 expression is highly effective. The additive effect observed with both constructs implies that residual Y-linked Cas9 expression enhances total cleavage efficiency within the X-poisoning system.

### qPCR Analysis of Y-Linked and Autosomal β-tubulin-Cas9 Expression

To compare transcriptional output between Y-linked and autosomal *βTub85D*-Cas9 constructs, Cas9 transcript levels were quantified in dissected testes and normalized to the housekeeping gene *RpL32*. Cas9 expression is shown as log-transformed fold change relative to wild type (Figure 3). Cas9 transcript abundance was strongly elevated in males carrying the autosomal construct (Aβ2-Cas9), whereas males carrying the Y-linked construct (Yβ2-Cas9) exhibited approximately 2,000-fold lower expression. Nonetheless, Cas9 transcripts in Yβ2-Cas9 testes remained significantly above wild-type background (*p* = 0.048, *t*-test), confirming that the *βTub85D* promoter retains limited transcriptional activity from the Y chromosome. **(Figure 3)**. These data confirm that *βTub85D*-driven transgenes are transcriptionally active but strongly downregulated when integrated into the Y chromosome, consistent with limited promoter accessibility in this chromatin context.

## Discussion

This study provides the first direct evidence that meiotic gene expression from the *D. melanogaster*□ Y chromosome is strongly repressed, though not completely silenced. While the spermatocyte-specific *βTub85D* promoter induced robust Cas9 activity from an autosomal locus, the same construct exhibited a roughly 2,000-fold reduction in transcript abundance when integrated into the Y chromosome and failed to induce measurable sex ratio distortion. These results indicate that *βTub85D*-driven transgenes retain transcriptional potential on the Y, □but are subject to substantial repression, consistent with pervasive heterochromatin and potential meiotic silencing.

Our findings parallel observations in *Anopheles gambiae*, where *β2-tubulin*-driven Y-linked reporters and SRD constructs show no detectable expression in the testes (Alcalay et al. 2021). However, *D. melanogaster*□ differs in that we observed detectable fluorescence in approximately two-thirds of Yβ2-Cas9 males and measurable Cas9 transcripts above background, suggesting that the *Drosophila* Y chromosome permits limited transcription from a meiotic promoter. Activity sufficient to enhance overall cleavage efficiency when combined with an autosomal copy, but not enough to independently drive SRD. The additive distortion observed in double-Cas9 males therefore suggests that even weak Y-linked expression can contribute functionally when paired with a strongly expressed autosomal allele.

The comparison between□ *An. gambiae*□ *and*□ *D. melanogaster*□□emphasizes both shared and species□specific features of Y□chromosome regulation. In mosquitoes, Y-linked transcription during meiosis appears almost completely suppressed, consistent with canonical MSCI. In contrast, the fly Y chromosome contains several large, actively transcribed fertility genes that form transcriptionally permissive nuclear loops in spermatocytes (Bonaccorsi and Lohe 1991; Piergentili 2007). Our results support a model in which the *Drosophila* Y can support low-level, spatially restricted transcription even from heterologous promoters, but not at the efficiency required for full functional output.

The repression of Y-linked *βTub85D*-Cas9 expression likely arises from the combined effects of heterochromatic spreading and incomplete escape from MSCI-like mechanisms. The *βTub85D* promoter, although active during meiosis, may lack the regulatory elements necessary to recruit the chromatin remodeling complexes that activate native Y fertility genes. Alternatively, integration–specific differences in chromatin accessibility could further constrain expression, despite the inclusion of insulating elements in our construct. The additive sex distortion observed in males carrying both autosomal and Y-linked Cas9 transgenes highlights the functional relevance of even weak Y-linked expression. This interaction mirrors the enhanced cleavage efficiency reported in multiplexed gRNA systems (Simoni et al. 2020) and suggests that Y-linked expression, while limited, could augment overall drive activity when paired with autosomal or X-linked components.

Collectively, these findings refine our understanding of Y-linked transcriptional capacity in *Drosophila* and emphasize that achieving robust meiotic expression from the Y chromosome will require more than classical promoter selection. Future research should prioritize identifying native Y-linked enhancers or promoters from genes known to be robustly transcribed during spermatogenesis in *D. melanogaster*, such as *kl-3* and *kl-5* (Vibranovski 2014) Additionally, exploiting epigenetic engineering tools to remodel local chromatin environments, or employing boundary elements to insulate transgenes from heterochromatin effects, represents a promising direction (Chiarella et al. 2020). The approaches developed here provide a framework for testing such strategies and, more broadly, for engineering Y-linked gene drives that exploit but also transcend the constraints of sex-chromosome biology.

## Methods

### Construct Design and Assembly

A ∼3.2 kb fragment encoding *β2-tubulin*-Cas9-T2A-eGFP was amplified from a previously validated construct (Haber et al. 2024) using primers BtubCas9_F and BtubCas9_R. The PCR product was cloned into an AscI-digested AByG backbone (Addgene plasmid #111083), which contains Y chromosome homology arms spanning nucleotide positions 666,900 to 667,710, as previously described by Buchman et al. (2019). Assembly was performed using Gibson cloning. The final construct consists of Y chromosome homology arms flanking *gypsy* and *CTCF* insulators, followed by the *βTub85D* promoter driving Cas9-T2A-eGFP, with′ a 3′ P10 terminator. Downstream a 3xP3 promoter drives expression of tdTomato for eye-specific fluorescent screening **(Figure 1A)**. The bacterial backbone included an ampicillin resistance gene and origin of replication.

### Embryo Microinjection and Line Establishment

The *βTub85D-Cas9-T2A-eGFP* construct was targeted to the *Drosophila melanogaster* Y chromosome using CRISPR/Cas9-mediated HDR. Microinjections were performed by Rainbow Transgenic Flies (Camarillo, CA, USA) into embryos from line 78782 (*nos*-Cas9; attP2, vermilion mutant background; BDSC #78782), following established protocols (Gamez et al. 2021). To enhance HDR efficiency, the injection mix included the donor plasmid, recombinant Cas9 protein (PNA Bio Inc., Newbury Park, CA), and in vitro transcribed sgRNAs targeting the Y-linked homology site. Surviving G□ adults were crossed to *w1118* flies, and progeny were screened for 3×P3-tdTomato fluorescence in the eyes. Transgenic males carrying the 3×P3-tdTomato marker were selected and backcrossed over multiple generations to *w1118* females to eliminate the endogenous *nanos*-Cas9 background.

Complete background replacement was confirmed by the absence of the *vermillion* eye phenotype. Chromosomal linkage of the transgene was verified by successive crosses between transgenic males and w1118 females. Across multiple generations, 3xP3-tdTomato expression was observed exclusively in male progeny and never detected in females, confirming stable Y chromosome integration **(Figure 1B)**. To assess transgene expression from the Y chromosome, testes were dissected from adult Yβ2Cas9 males in 1× PBS and examined under an Olympus SZX16 stereomicroscope. GFP fluorescence was evaluated in freshly dissected samples, and images were captured using an EVOS M7000 Imaging System (Thermo Fisher Scientific). As a positive control, testes were similarly dissected and imaged from males carrying an autosomal insertion of the same *βTub85D-Cas9* construct (Aβ2Cas9) **(Figure 1C)**.

### Fly Husbandry

All *Drosophila melanogaster* stocks and genetic crosses were maintained under standard laboratory conditions at 25□°C with a 12:12-hour light–dark cycle. Flies were reared on standard cornmeal-agar-yeast food and transferred to fresh food vials every 2-3 weeks to maintain colony health and reproductive viability. Routine handling and phenotyping were performed under CO□ anesthesia.

Transgenic individuals were screened for fluorescence using a stereomicroscope equipped with the appropriate filter sets. The autosomal Cas9 source, integrated on the third chromosome, marked by an *Opie2*-driven DsRed marker, whereas the *RpS6*-targeting *gRNA* construct, inserted on the second chromosome, marked by an *3xP3*-eGFP fluorescent cassette, as previously described by Haber et al. (2024). The Y-linked Cas9 construct, generated following the approach of Buchman and Akbari (2019), was marked with a 3×P3-tdTomato (RFP) reporter to facilitate identification of male carriers.

### Sex ratio distortion assays

To evaluate sex ratio distortion (SRD) efficiency, we performed a series of genetic crosses combining different Cas9 configurations with the *RpS6_2* gRNA line, which targets an X-linked haplolethal ribosomal protein gene. For the Y-linked construct, males carrying the *βTub85D*-Cas9 transgene on the Y chromosome were crossed with *RpS6_2* gRNA females to generate F1 trans-heterozygous males (Yβ2Cas9*/RpS6_2*). For the autosomal Cas9 line, virgin females carrying the *βTub85D*-Cas9 construct were crossed with *RpS6_2* gRNA males to produce F1 trans-heterozygous males (Aβ2Cas9*/RpS6_2*). For experiments combining both constructs, Btub_Y males were crossed with virgin females carrying the autosomal *βTub85D*n-Cas9 transgene to generate double-transgenic males carrying both Y-linked and autosomal Cas9 insertions. Transgene presence was confirmed by fluorescence screening. The autosomal Cas9 construct carried an Opie2-DsRed marker, and the Y-linked construct carried 3×P3-tdTomato, both were detected as RFP fluorescence. For single-Cas9 genotypes, RFP signal confirmed transgene presence. These males were crossed to *RpS6_2* gRNA flies, which carried a 3×P3-GFP marker, to generate trans-heterozygotes (A+Yβ2Cas9/*RpS6_2*). In all cross types, the resulting F1 trans-heterozygous males were subsequently backcrossed to wild-type (*w1118*) females, and the sex ratio of their progeny was recorded. As negative controls, crosses were performed without the *RpS6_2* gRNA line. Males carrying only the Cas9 transgene (Yβ2Cas9*/+*, Aβ2Cas9*/+*, or A+Yβ2Cas9*/+*) were crossed to wild-type females to assess the baseline sex ratio in the absence of X-poisoning. All crosses were conducted in vials containing five males and five females to ensure consistency across experimental groups.

### qPCR for Cas9 expression

Total RNA was extracted from the testes of adult males carrying the Y-linked *βTub85D*-Cas9 construct (Yβ2Cas9), the autosomal construct (Aβ2Cas9), and wild-type controls. Three biological replicates were analyzed for each genotype, each consisting of 20 pooled testes. RNA was isolated using the MicroRNA Kit (Zymo Research, Irvine, CA, USA) according to the manufacturer’s protocol. Complementary DNA (cDNA) was synthesized from 1□μg of total RNA using the High-Capacity cDNA Reverse Transcription Kit (Thermo Fisher Scientific, Waltham, MA, USA) with random hexamer primers. Quantitative PCR (qPCR) was performed using SYBR Green chemistry on a QuantStudio 3 Real-Time PCR System (Applied Biosystems). Each reaction was run in technical triplicate in a 20□μL total volume. Cas9 transcript levels were amplified using the primer pair Cas9_qPCR_F and Cas9_qPCR_R and normalized to the housekeeping gene RpL32, amplified with RpL32_qPCR_F and RpL32_qPCR_R. Relative expression levels were calculated using the ΔΔCt method, and expression from each Cas9 genotype was compared to wild-type controls.

## Author Contributions

YA and PAP conceived and designed the study. YA, CZ generated transgenic constructs and strains and performed all fly experiments. EY helped in genomic and bioinformatic analysis. EB and EY designed and performed the quantitative RT-PCR analysis. YA and PAP supervised the project, secured funding, and wrote the manuscript together with CZ with input from all authors. All authors read and approved the final manuscript.

## Acknowledgements

We thank Eyal Schechter for his early guidance and for sharing his extensive knowledge of Drosophila biology and genetics. We are also grateful to Flavia Krsticevic, Daniella An Haber, Nikolai Windbichler, and Omar Akbari for valuable discussions and critical feedback throughout the project. We thank Moshe Elbaz for his support in setting up this work. Fly icons (fly_female_adult_from_a_lateral_view and fly_male_adult_from_a_lateral_view) by DBCLS and licensed under CC-BY 4.0 unported were downloaded from Bioicons.com.

## Ethics

All insect work was performed in facilities maintaining Arthropod Containment Level 2. This work received Institutional Approval and relevant authorizations from the Israel Ministry of Environmental Protection and Ministry of Agriculture (#1031/21).

## References

Alcalay, Yehonatan, Silke Fuchs, Roberto Galizi, et al. 2021. “The Potential for a Released Autosomal X-Shredder Becoming a Driving-Y Chromosome and Invasively Suppressing Wild Populations of Malaria Mosquitoes.” Frontiers in Bioengineering and Biotechnology 9 (December). 10.3389/fbioe.2021.752253.

Alphey, Luke. 2014. “Genetic Control of Mosquitoes.” Annual Review of Entomology 59 (Volume 59, 2014): 205–24. 10.1146/annurev-ento-011613-162002.

Anderson, James T, Steven Henikoff, and Kami Ahmad. 2023. “Chromosome-Specific Maturation of the Epigenome in the Drosophila Male Germline.” eLife 12 (November): RP89373. 10.7554/eLife.89373.

Bonaccorsi, S, and A Lohe. 1991. “Fine Mapping of Satellite DNA Sequences along the Y Chromosome of Drosophila Melanogaster: Relationships between Satellite Sequences and Fertility Factors.” Genetics 129 (1): 177–89. 10.1093/genetics/129.1.177.

Buchman, A., and O. S. Akbari. 2019. “Site-Specific Transgenesis of the Drosophila Melanogaster Y-Chromosome Using CRISPR/Cas9.” Insect Molecular Biology 28 (1): 65– 73. 10.1111/imb.12528.

Burt, Austin, and Anne Deredec. 2018. “Self-Limiting Population Genetic Control with Sex-Linked Genome Editors.” Proceedings. Biological Sciences 285 (1883): 20180776. 10.1098/rspb.2018.0776.

Carvalho, A. Bernardo. 2002. “Origin and Evolution of the Drosophila Y Chromosome.” Current Opinion in Genetics & Development 12 (6): 664–68. 10.1016/S0959-437X(02)00356-8.

Compton, Austin, and Zhijian Tu. 2022. “Natural and Engineered Sex Ratio Distortion in Insects.” Frontiers in Ecology and Evolution 10 (June). 10.3389/fevo.2022.884159.

Deredec, Anne, H. Charles J. Godfray, and Austin Burt. 2011. “Requirements for Effective Malaria Control with Homing Endonuclease Genes.” Proceedings of the National Academy of Sciences 108 (43): E874–80. 10.1073/pnas.1110717108.

Fasulo, Barbara, Angela Meccariello, Maya Morgan, Carl Borufka, Philippos Aris Papathanos, and Nikolai Windbichler. 2020. “A Fly Model Establishes Distinct Mechanisms for Synthetic CRISPR/Cas9 Sex Distorters.” PLOS Genetics 16 (3): e1008647. 10.1371/journal.pgen.1008647.

Fuller, M.T. 1993. “Spermatogenesis.” In The Development of Drosophila Melanogaster, vol. 1. Cold Spring Harbor Laboratory Press.

Galizi, Roberto, Andrew Hammond, Kyros Kyrou, et al. 2016. “A CRISPR-Cas9 Sex-Ratio Distortion System for Genetic Control.” Scientific Reports 6 (1): 31139. 10.1038/srep31139.

Gamez, Stephanie, Duverney Chaverra-Rodriguez, Anna Buchman, et al. 2021. “Exploiting a Y Chromosome-Linked Cas9 for Sex Selection and Gene Drive.” Nature Communications 12 (1): 7202. 10.1038/s41467-021-27333-1.

Haber, Daniella An, Yael Arien, Lee Benjamin Lamdan, et al. 2024. “Targeting Mosquito X-Chromosomes Reveals Complex Transmission Dynamics of Sex Ratio Distorting Gene Drives.” Nature Communications 15 (1): 4983. 10.1038/s41467-024-49387-7.

Hense, Winfried, John F. Baines, and John Parsch. 2007. “X Chromosome Inactivation during Drosophila Spermatogenesis.” PLOS Biology 5 (10): e273. 10.1371/journal.pbio.0050273.

Kemkemer, C., A. Catalán, and J. Parsch. 2014. “‘Escaping’ the X Chromosome Leads to Increased Gene Expression in the Male Germline of Drosophila Melanogaster.” Heredity 112 (2): 149–55. 10.1038/hdy.2013.86.

Kemkemer, Claus, Winfried Hense, and John Parsch. 2011. “Fine-Scale Analysis of X Chromosome Inactivation in the Male Germ Line of Drosophila Melanogaster.” Molecular Biology and Evolution 28 (5): 1561–63. 10.1093/molbev/msq355.

Krzywinski, Jaroslaw, Mathew A. Chrystal, and Nora J. Besansky. 2006. “Gene Finding on the Y: Fruitful Strategy in Drosophila Does Not Deliver in Anopheles.” Genetica 126 (3): 369–75. 10.1007/s10709-005-1985-3.

Landeen, Emily L., Christina A. Muirhead, Lori Wright, Colin D. Meiklejohn, and Daven C. Presgraves. 2016. “Sex Chromosome-Wide Transcriptional Suppression and Compensatory Cis-Regulatory Evolution Mediate Gene Expression in the Drosophila Male Germline.” PLOS Biology 14 (7): e1002499. 10.1371/journal.pbio.1002499.

Mahadevaraju, Sharvani, Justin M. Fear, Miriam Akeju, et al. 2021. “Dynamic Sex Chromosome Expression in Drosophila Male Germ Cells.” Nature Communications 12 (1): 892. 10.1038/s41467-021-20897-y.

Meccariello, Angela, Flavia Krsticevic, Rita Colonna, et al. 2021. “Engineered Sex Ratio Distortion by X-Shredding in the Global Agricultural Pest Ceratitis Capitata.” BMC Biology 19 (1): 78. 10.1186/s12915-021-01010-7.

Meiklejohn, Colin D., Emily L. Landeen, Jodi M. Cook, Sarah B. Kingan, and Daven C. Presgraves. 2011. “Sex Chromosome-Specific Regulation in the Drosophila Male Germline But Little Evidence for Chromosomal Dosage Compensation or Meiotic Inactivation.” PLOS Biology 9 (8): e1001126. 10.1371/journal.pbio.1001126.

Piergentili, Roberto. 2007. “Evolutionary Conservation of Lampbrush-like Loops in Drosophilids.” BMC Cell Biology 8 (1): 35. 10.1186/1471-2121-8-35.

Simoni, Alekos, Andrew M. Hammond, Andrea K. Beaghton, et al. 2020. “A Male-Biased Sex-Distorter Gene Drive for the Human Malaria Vector Anopheles Gambiae.” Nature Biotechnology 38 (9): 1054–60. 10.1038/s41587-020-0508-1.

Vibranovski, Maria D. 2014. “Meiotic Sex Chromosome Inactivation in Drosophila.” Journal of Genomics 2 (June): 104–17. 10.7150/jgen.8178.

Vibranovski, Maria D., Yong Zhang, and Manyuan Long. 2009. “General Gene Movement off the X Chromosome in the Drosophila Genus.” Genome Research 19 (5): 897–903. 10.1101/gr.088609.108.

Witt, Evan, Zhantao Shao, Chun Hu, Henry M. Krause, and Li Zhao. 2021. “Single-Cell RNA-Sequencing Reveals Pre-Meiotic X-Chromosome Dosage Compensation in Drosophila Testis.” PLOS Genetics 17 (8): e1009728. 10.1371/journal.pgen.1009728.

