## Supplementary Table 1 and 2 for "Developing Y chromosome sex ratio distorters in the model insect *Drosophila melanogaster*"

**Supplementary Table 1. Summary of embryo injections for Y-linked transgene integration**

| Injection Round | Injected Eggs | Surviving Larvae (%) | RFP+ Males Recovered | Outcome Summary |
|---|---|---|---|---|
| 1 | 245 | 155 (63%) | 0 | No transformants |
| 2 | 255 | 150 (59%) | 0 | No transformants |
| 3 | 150 | 91 (60%) | 4 | 3 sterile; 1 line with Y-linked insertion confirmed |

**Supplementary Table 2. Primer sequences used for cloning and gene expression analysis**

| Primer name | Sequence (5′ → 3′) | Purpose |
|---|---|---|
| BtubCas9_F | **GTCAATATCAATCGTATCATCTGGTCG**AGCACGGGACGTGCGACG | Amplification of *β2Tubulin*-Cas9-T2A-eGFP for Cloning (Gibson) (forward) |
| BtubCas9_R | **GTTAACTCGAATCGCTATCCAAGC**TACGGTAGCAGAGACTTGGTCTCCTAAATTCGAGCTCGCCCGGG | Amplification of *β2Tubulin*-Cas9-T2A-eGFP for Cloning (Gibson) (reverse) |
| Cas9_qPCR_F | CCTACAACAAGCACCGGGAT | qPCR (forward) |
| Cas9_qPCR_R | TACCTCTTCCGGTCGATGGT | qPCR (reverse) |
| RpL32_qPCR_F | GACAATCTCCTTGCGCTTCT | qPCR reference (forward) |
| RpL32_qPCR_R | ATCGGTTACGGATCGAACAA | qPCR reference (reverse) |
